# Development and analytical validation of a finite element model of fluid transport through osteochondral tissue

**DOI:** 10.1101/2020.10.26.356188

**Authors:** Brady D. Hislop, Chelsea M. Heveran, Ronald K. June

## Abstract

Fluid transport between cartilage and bone is critical to joint health. The objective of this study was to develop and analytically validate a finite element model of osteochondral tissue capable of modeling cartilage-bone fluid transport. A biphasic viscoelastic model using an ellipsoidal fiber distribution was created with three distinct layers of cartilage (superficial zone, middle zone, and deep zone) along with a layer of subchondral bone. For stress-relaxation in unconfined compression, our results for compressive stress, radial stress, effective fluid pressure, and elastic recoil were compared with established biphasic analytical solutions. Our model also shows the development of fluid pressure gradients at the cartilage-bone interface during loading. Fluid pressure gradients developed at the cartilage-bone interface with consistently higher pressures in cartilage following initial loading to 10% strain, followed by convergence towards equal pressures in cartilage and bone during the 400s relaxation period. These results provide additional evidence that fluid is transported between cartilage and bone during loading and improves upon estimates of the magnitude of this effect through incorporating a realistic distribution and estimate of the collagen ultrastructure. Understanding fluid transport between cartilage and bone may be key to new insights about the mechanical and biological environment of both tissues in health and disease.

## Introduction

Joint injuries frequently progress to debilitating osteoarthritis (OA), a leading cause of disability worldwide. For example, 40% of patients who experience a traumatic joint injury will develop post-traumatic OA (PTOA) within 10 years^36^. Cartilage and subchondral bone serve as the key structural components of our joints, with cartilage serving as the load-bearing tissue and subchondral bone providing structural support. Cartilage is a smooth avascular tissue that relies on its surrounding environment for nourishment. The current paradigm for joint fluid transport is that cartilage receives its nutrients through the synovial fluid and lymph nodes^28^. By contrast, bone is vascularized, which allows the rapid transport of nutrients as well as cytokines^10^.

It is possible that fluid is transported directly between bone and cartilage. This transport could potentially greatly increase the nutrients and signaling molecules available to chondrocytes compared with transport from the synovial fluid and lymph. Imaging studies of the cartilage-bone interface show the transport of solutes *in vivo* between the subchondral bone and calcified cartilage^29,30^. Further, a poroelastic finite element model demonstrated that joint loading and unloading promotes fluid movement between cartilage and bone^35^. However, specific gaps in knowledge still remain about how fluid is transported between these tissues.

Specifically, the underlying mechanics responsible for fluid transport are still heavily debated with differing results between models using either a viscoelastic solid matrix or poroelastic approach^8,11,32^. Prior models of cartilage mechanics either neglect collagen fibers or model collagen fibers in a simplified manner to reduce model complexity^12,32^. Stender *et al* approximate collagen ultrastructure with an anisotropic fiber distribution. Here, we improve upon this estimate by using an ellipsoidal fiber distribution^3,4^. The ellipsoidal fiber distribution mimics the Benninghoff arcade geometry seen in articular cartilage with zone-dependent fiber orientation^5^. Incorporating this fiber distribution into an osteochondral fluid transport model is important because the orientation of collagen fibers influences the mechanical environment, and potentially fluid transport properties, of cartilage^3,4,28^. Additionally in this study, we use a viscoelastic solid phase. Many previous studies assume poroelasticity^13,32^ of the solid phase, but this assumption is challenged by observations of cartilage viscoelastic behavior in pure shear^11,15^. A biphasic viscoelastic model was chosen with the goals of capturing the viscoelastic responses of the extracellular matrix and collagen fibers as well as the interactions between the solid phase and water.

The overall objective of this study is to develop a finite element model of cartilage-bone fluid transport that is analytically validated with established biphasic solutions for elastic recoil, fluid pressure and radial size effects^2^. We hypothesize that fluid is transported between cartilage and bone during physiological loading/unloading and is driven by cartilage elastic recoil. Therefore, the three goals of this study are: (i) develop a finite element model of fluid transport in osteochondral tissue mimicking an unconfined compression experiment at physiologically relevant loads (ii) validate the finite element model against established analytical solutions; and (iii) quantify fluid pressure gradients at the cartilage-bone interface.

## Materials and Methods

### Model Geometry

Osteochondral cylinders (1-2mm diameter and 4mm height) were modeled in FEBio^21^ using quarter symmetry (Figure 1). Boundary conditions simulated unconfined compression with impermeable boundaries and fixed displacements perpendicular to each side. To minimize nonlinear effects and computational complexity, a single body geometry was selected with hexahedral and penta mesh elements. Spatially dependent cartilage and bone material models were defined according to a 50-15-25-10 percentage split of bone, deep, middle, and superficial cartilage zones, respectively, using an in-house MATLAB code (Figure 1).

**Figure 1.**
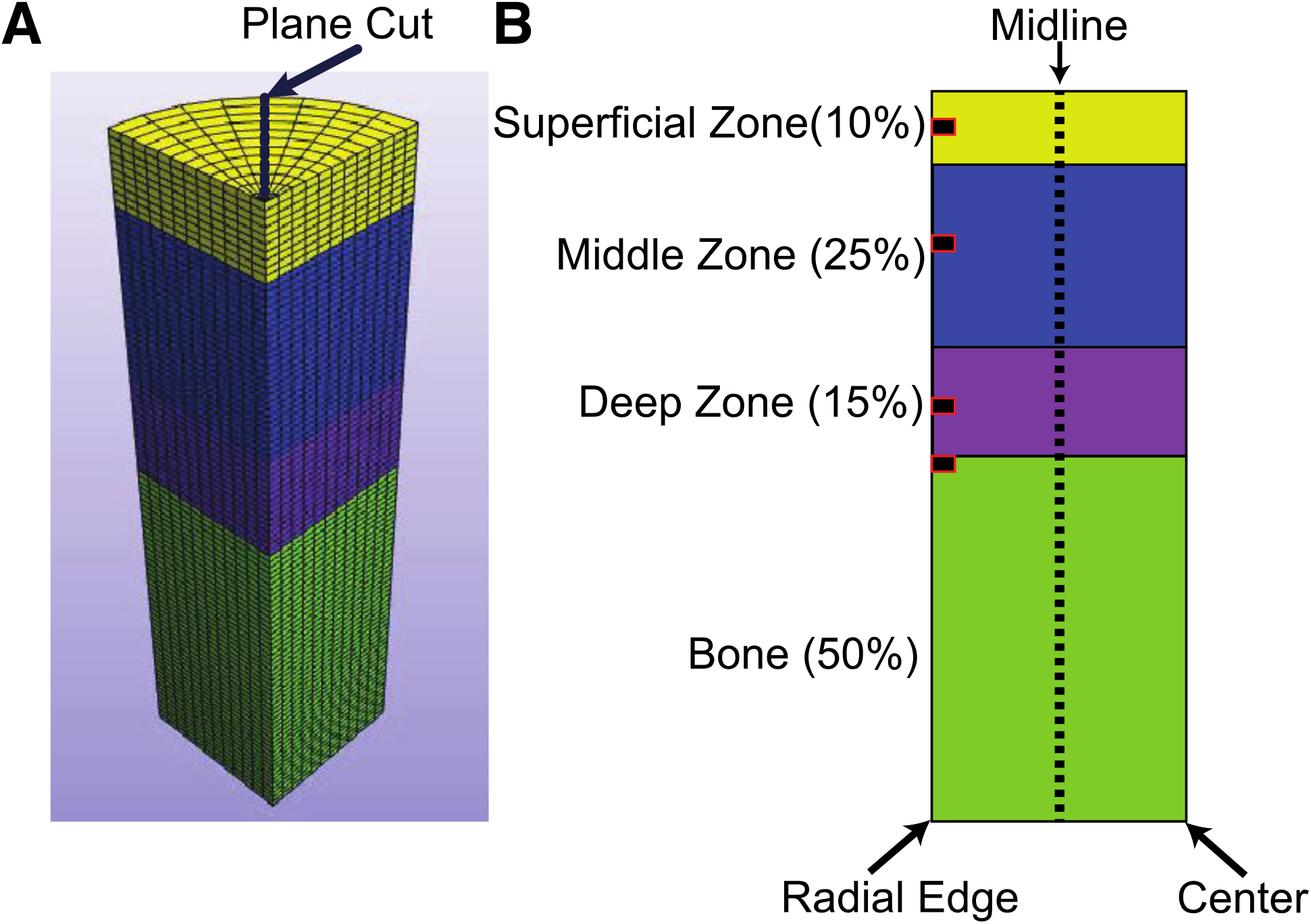
Finite element model of osteochondral tissue A) Mesh showing plane cut for evaluation; B) Schematic of cartilage layers and bone showing location of elements that are plotted in Figures 2-5.

### Cartilage constitutive model and material properties

A viscoelastic neo-Hookean ellipsoidal fiber distribution model^3,4,28^ was chosen for cartilage. Such modeling combines the tensile properties of collagen fibers and the biphasic viscoelastic response of the remaining matrix components and their bound water. The viscoelastic neo-Hookean model of the remaining matrix was applied to each layer of the tissue (v=0.499, relaxation time = 120s)^23^. A depth-dependent elastic modulus of the tissue was derived and applied to each layer of cartilage, using data from Walhquist *et al.*^34^:

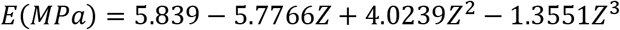

where Z is the depth from the cartilage-bone interface. In addition, an ellipsoidal fiber distribution^3,4^ was applied to each layer of cartilage independently with superficial zone fibers parallel to the superficial surface *(β =* (2.5,2.5,2.5), *ζ =* (4,4,2)), middle zone fibers uniformly distributed *(β =* (3.5,3.5,3.5), *ζ =* (4,4,4)), and deep zone fibers perpendicular to the superficial surface (β = (2.5,2.5,2.5), ζ = (2,2,4)), with *β* defining the elastic modulus along each axis and *ζ* defining the spatial distribution of fibers. The biphasic nature of the tissue was accounted for through the implementation of the Holmes-Mow permeability function^18^, modeling the strain-dependent permeability of cartilage (*k_o_* = 6.2*e*^−16^,*M* = 0.4, *α* = 2.2), where *k_o_* defines the initial permeability, *α* defines the rate at which permeability approaches zero, and M is constant for the exponential fit.

### Bone constitutive model and material properties

Previous computational studies of the osteochondral tissues have modeled bone as a rigid body given the large difference in modulus between cartilage and bone (bone ~1000x greater) between cartilage and bone. However, rigid body mechanics cannot be adapted for biphasic materials. Therefore, the bone was modeled as a biphasic isotropic elastic material (E= 17GPa, v= 0.29, φ= 5.5%)^1,35^ throughout the geometry. Isotropic elastic models are considered effective for modeling the mechanical properties of bone^14,15^ while allowing for the implementation of biphasic properties. Given the relatively low levels of bone deformation during the simulated loading, the bone permeability was set as spatially constant (1e^−17^)^35^.

### Verification and model studies

Stress-relaxation was simulated using two simulation steps: a ramped compression step followed by a hold at 10% strain for 400s. Vertical and radial mesh convergence studies were performed to determine the appropriate mesh size for this model. Effective fluid pressures of each mesh were studied to determine the appropriate mesh size, using a threshold of < 5% change to determine convergence. Additionally, a radial size study was performed to determine the effects of various radii on the physical parameters of the model. Each model’s results were analyzed for vertical and radial stress relaxation, radial elastic recoil, effective fluid pressure relaxation, and differences in these physical parameters for the various radii to compare against validated analytical solutions^2^. Analytical predictions included Poisson’s-ratio dependent elastic recoil and increasing radial elastic recoil with decreasing Poisson’s ratio. Further analytical predictions included increased magnitude of compressive stress with increased radius. All results during this study were analyzed in PostView^22^ a post-processing software developed to support FEBio. All simulations were run on the Hyalite Cluster at Montana State University^35^ using 8 cores and 4GB RAM per core to achieve rapid solutions (~4 hrs).

## Results

### Mesh Convergence

Evaluation of the effective fluid pressure across three vertical mesh sizes (100, 200, 300 layers per z-stack) revealed a < 5% increase in effective fluid pressure between 100 and 200 layers, and 200 and 300 layers (Sup. Figure 1B). Whereas, in the radial mesh study (8, 12, 16 radial slices) we saw increases in effective fluid pressure of 11.1% and 4.7% between 8 and 12, 12 and 16, respectively (Sup. Figure 1A). Given these results, we built our mesh with 8 theta segments, 12 radial slices, and 100-vertical layers.

#### Compressive stress relaxation

Compressive stress relaxation in the z-direction was observed within both cartilage and bone elements. Detailed inspection along the radial edge revealed z-direction relaxation throughout each layer of cartilage (superficial, middle, and deep; Figure 2A-F). While further evaluation revealed stress relaxation along the midline and center of the tissue for each layer (supplemental movie 1). Additionally, stress relaxation was observed throughout the bone, with decreasing levels of compressive stress with depth from the cartilage-bone interface.

**Figure 2.**
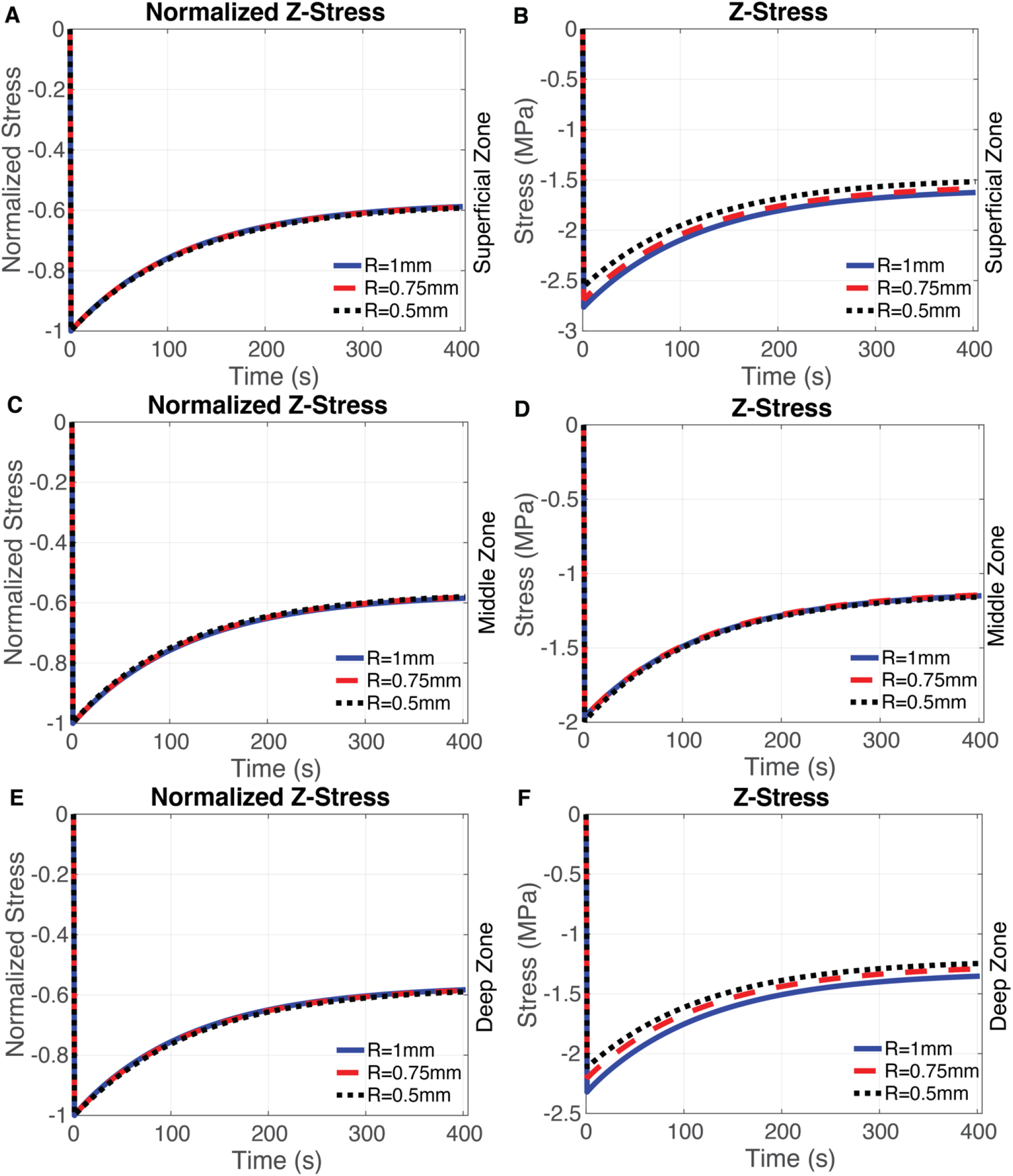
Compressive stress as a function of time for three radii. Normalized to peak compressive stress (left column) and absolute stress (right column) at location of interest and compressive stress at mid-depth of the superficial-zone (A-B), middle-zone (C-D), and deepzone (E-F).

Radial mesh studies revealed identical relaxation profiles across each radial mesh size (1, 0.75 and 0.5 mm). We also found that levels of compressive stress had a direct relationship with decreases in radial mesh size. Suggesting that our mesh size is appropriate for modeling the expected decreases in compressive stress with reductions in the radius of the model. Additionally, using a stretched exponential function^11^ we found a relaxation time constant of 969.5 seconds.

#### Radial stress relaxation

Radial stress relaxation at the radial edge (r=R) of cartilage was observed through each layer of the tissue. The middle zone of cartilage showed negative overall radial stress (Figure 3C-D), while the superficial zone showed an increase in radial stress from 0.5 mm to 1 mm (Figure 3F). Similar relaxation was seen at the radial edge of bone throughout most of the depth. However, near the bottom of the bone, a wave response was observed following the initial load.

**Figure 3.**
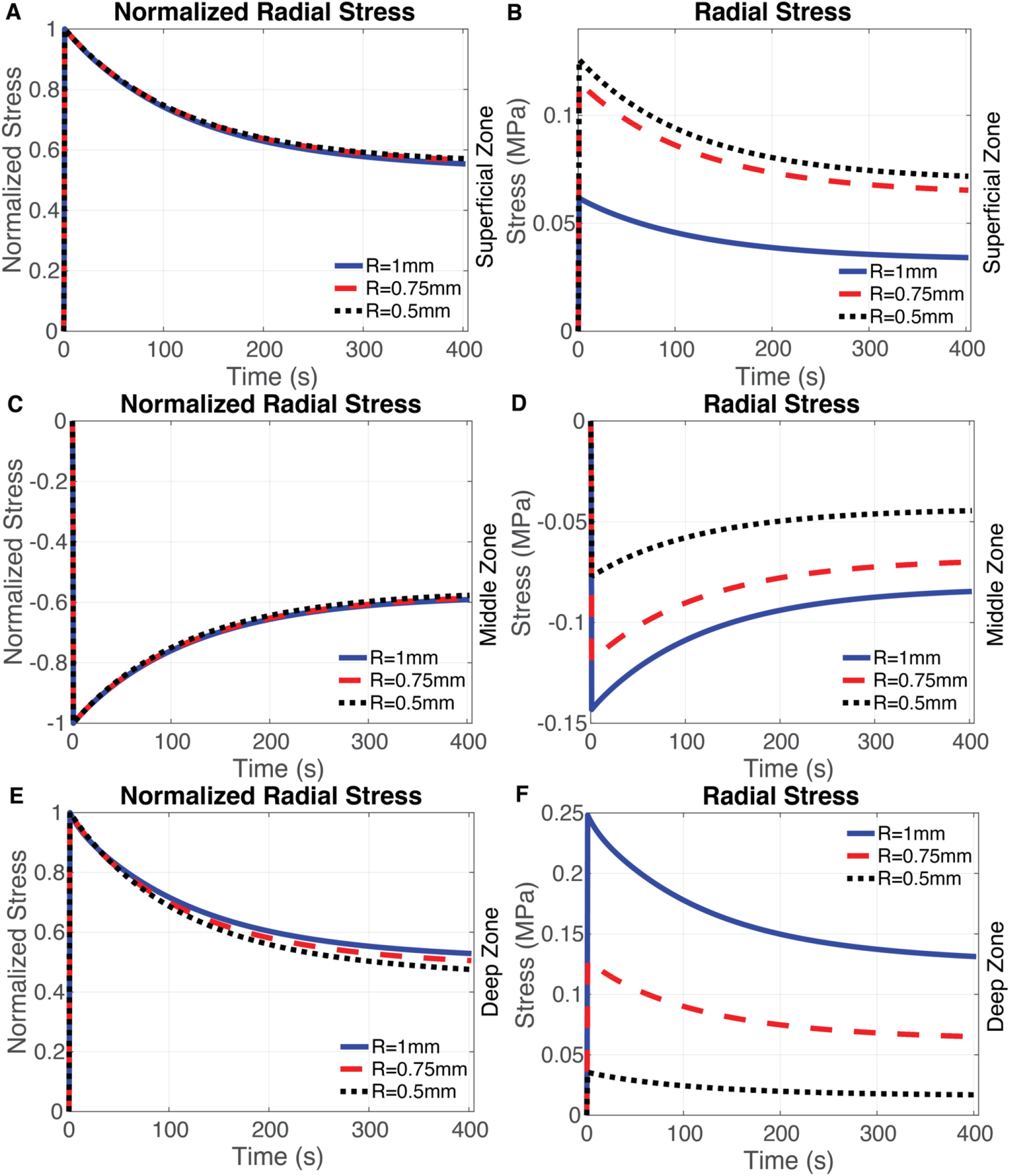
Radial stress as a function of time for three radii. Stress normalized to the peak radial stress at location of interest (left column) and absolute radial stress (right column) at the middepth of the superficial-zone(A-B), middle-zone (C-D), and deep-zone (E-F).

Slight deviations in normalized relaxation profiles were observed between each radial mesh size with increases in relaxation time as the radius decreased in the deep zone of cartilage (Figure 3A). Similar to the effective pressure and compressive stress radial stress has a direct relationship with decreases in radial mesh size.

#### Effective Fluid pressure relaxation

Effective fluid pressure relaxation was observed throughout cartilage and bone following the initial loading of cartilage. Analysis of the radial edge revealed fluid pressure relaxation for each layer of cartilage (Figure 4A-F) and radial mesh size. In addition, analysis of the midline and center of cartilage revealed similar patterns of fluid pressure relaxation. Likewise, analysis of bone at the radial edge and along the midline revealed fluid pressure relaxation throughout the bone. Additional analysis showed that pressure relaxation along the center of bone transitioned from negative to positive with depth from the cartilage-bone interface.

**Figure 4.**
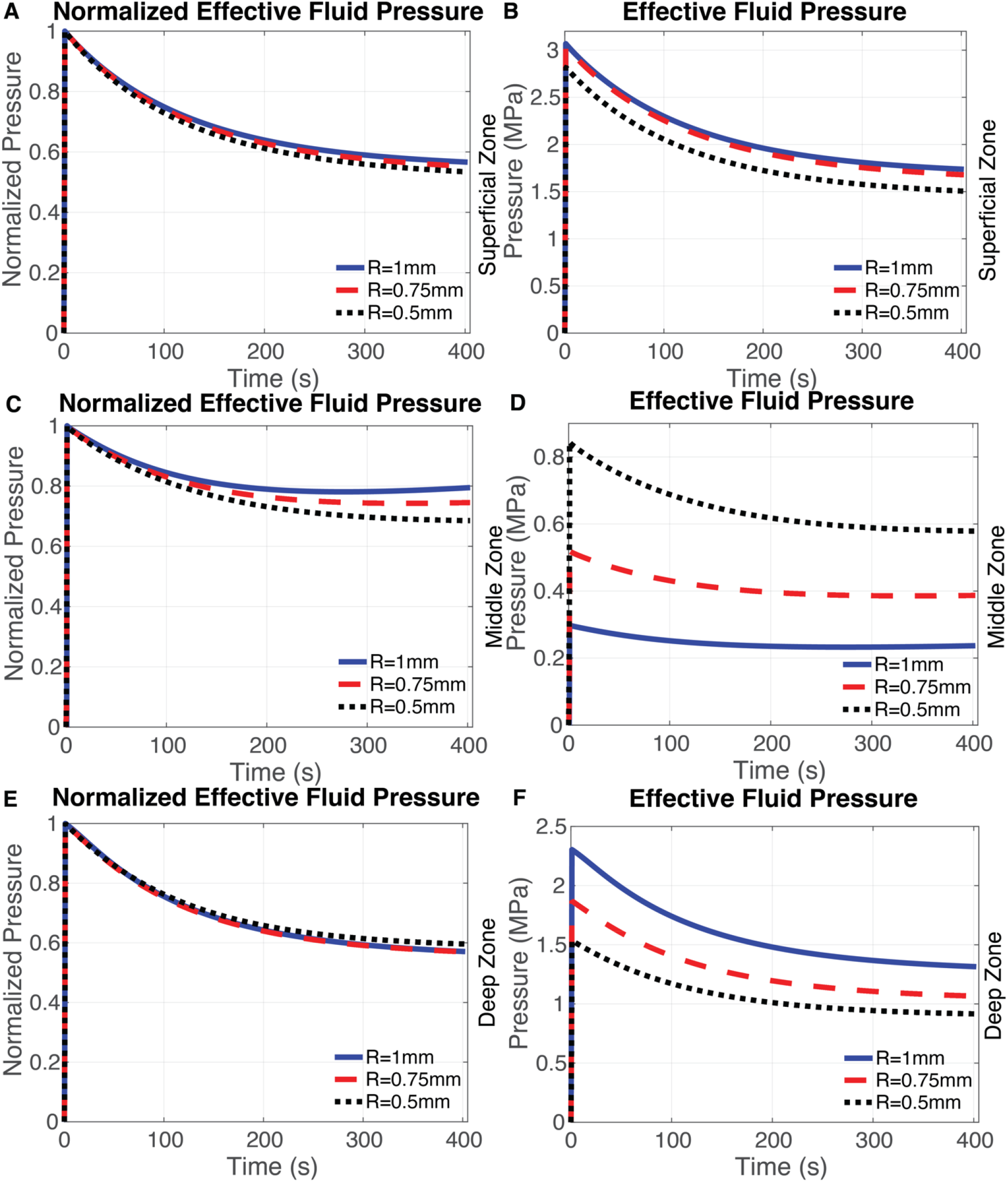
Effective fluid pressure in cartilage as a function of time for three radii. Fluid pressure normalized to peak effective fluid pressure (left column) and absolute effective fluid pressure (right column) at locations of interest at mid-depth of the superficial-zone (A-B), middle-zone (CD), and deep-zone (E-F).

Fluid pressure relaxation profiles were found to be similar for each of the radial mesh sizes (1, 0.75, 0.5mm) studied, with R = 0.5 mm having the highest relative relaxation after 400s (Figure 4A). Additionally, consistent with the results seen for compressive stress the effective fluid pressure has a direct relationship with decreases in radial mesh size.

#### Elastic Recoil

Elastic recoil of cartilage (v=0.499) at the radial edge was nonexistent (Figure 5A-C), whereas bone (v=0.29) (Figure 5D) underwent elastic recoil consistent with analytical solutions for biphasic materials^2^. We also found that levels of radial displacement decreased concurrently with a radial size consistent with the changes in compressive stress levels. Elastic recoil for cartilage was consistent across each of the radial mesh sizes and cartilage layers (Figure 5A-C). Conversely, bone recoil was the same across the 0.75- and 1-mm with changes in levels of recoil occurring only for the 0.5 mm model.

**Figure 5.**
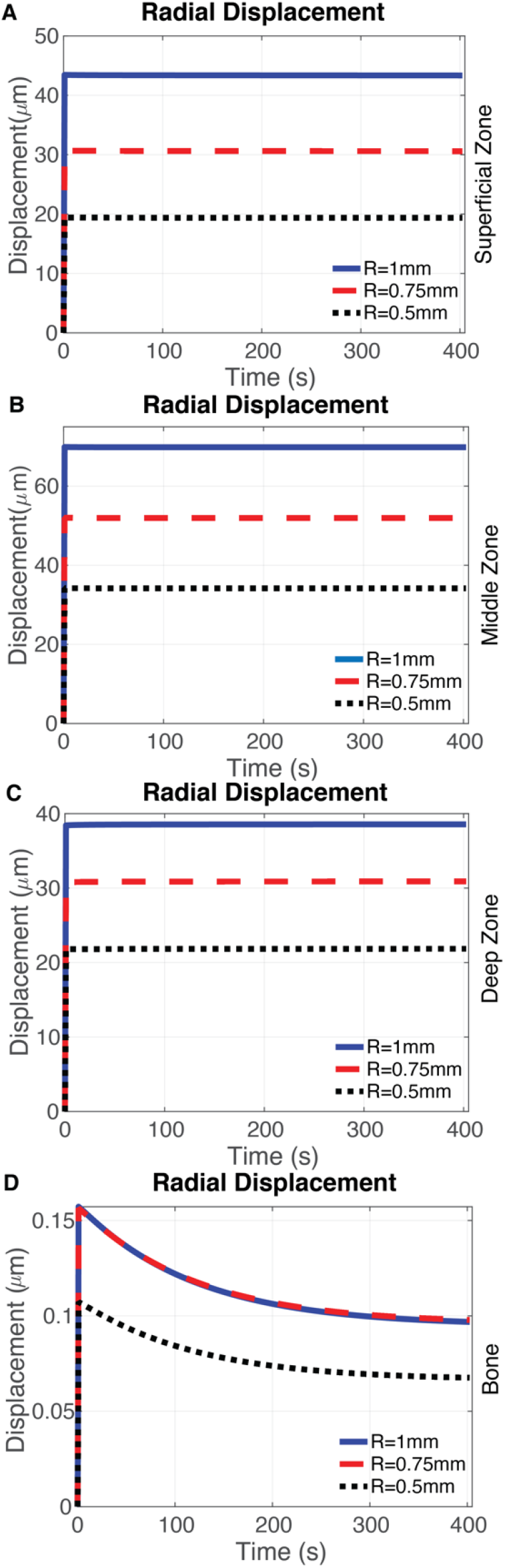
A-C: Radial Displacement of cartilage at mid-depth of superficial-zone, middle-zone and deep-zone for three radial mesh sizes; D: Radial Displacement of subchondral bone directly below cartilage for three radial mesh sizes.

#### Fluid Pressure Gradients

Cartilage fluid pressures at the cartilage-bone interface were consistently higher than bone immediately following the initial loading period (t=1 second) (Figure 6A). We also found that fluid pressure gradients decreased after the 400 second relaxation period (t = 402.5 seconds) (Figure 6B).

**Figure 6.**
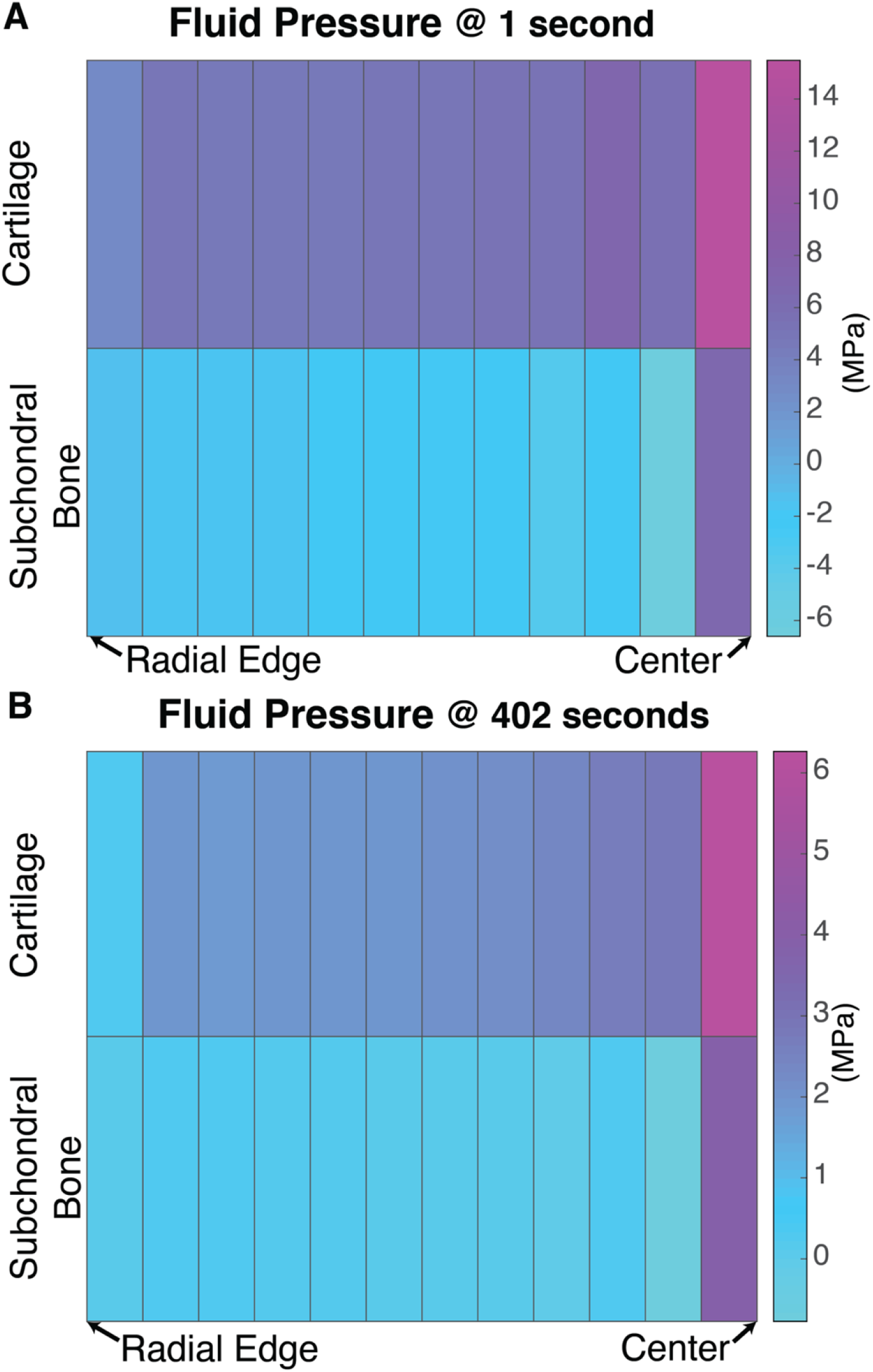
Mesh elements at the cartilage-bone interface A): Fluid pressures at cartilage-bone interface after initial load; B) Fluid pressure at cartilage-bone interface after 400 seconds of relaxation

## Discussion

The purpose of this study was to investigate fluid pressure gradients and cartilage elastic recoil in fluid transport across the cartilage-bone interface during unconfined compression experiments. Model results were consistent with analytical results for biphasic materials^2^, including axial and radial stress relaxation, elastic recoil, and effective fluid pressure relaxation. Results for the radial stresses in the superficial and middle zones of cartilage can be explained by the collagen fiber orientation in each region. Greater volume fraction of collagen fibers parallel to the cartilage surface (including many in the radial direction) in the superficial zone lead to lower radial stresses (Figure 3D) as the sample radius increases. By contrast, the random distribution of fibers in the middle zone and increased modulus (3.5 MPa v. 2.5 MPa) leads to lower stress magnitudes and overall negative radial stresses. This model also captures the Poisson’s ratio-dependent characteristics of elastic recoil for bone and cartilage. Further, the cartilage relaxation time of 969 seconds was within 20% of a previous experimental study^31^.

Modeling results at the cartilage-bone interface demonstrate that unconfined compression results in higher fluid pressures in the cartilage than bone. These results suggest that a fluid pressure gradient develops from cartilage to bone during loading (Figure 6A). Our results agree with Stender *et al* in that fluid is transported between cartilage and bone and this transport is driven by the development of fluid pressure gradients at the cartilage-bone interface. Stender *et al* presented models of both unconfined compression and spherical indentation toward understanding the influence of permeability on cartilage-bone fluid transport. Our model is also in agreement with these previous results on the direction of the fluid transport during loading, although our model predicts much lower levels of transport (~500 fold decreased flux) from cartilage to bone. This discrepancy may result from our use of an ellipsoidal fiber distribution and the previous use of an anisotropic distribution of collagen fibers. Additionally, our use of a viscoelastic model versus the previous use of a poroelastic model may also be important. Our model assumes that the matrix and fibers react in a time-dependent manner (viscoelastic) that is independent of the fluid movement, whereas the previous model assumes that the solid matrix and collagen fibers relax due to the movement of fluid inside the matrix (poroelastic). A limitation of both viscoelastic and poroelastic models is that neither fully describes cartilage time-dependent material properties. However, since our viscoelastic model and previous poroelastic models both produce the result of higher fluid pressure in cartilage than bone in loading, this fluid transport behavior appears robust to a range of assumptions about cartilage mechanics.

Our results challenge the conventional understanding that fluid transport predominantly occurs between cartilage and synovial fluid. Fluid transport from subchondral bone to cartilage *(i.e.,* during unloading) and within cartilage may impact the health and viability of chondrocytes. The transfer of nutrients and cytokines from subchondral bone could provide biochemical energy and chemical signals to chondrocytes. In this way, physiological loading could support chondrocyte health and viability by supplementing the nutrient transfer from synovial fluid. An additional interpretation of our results is that bone may act as an ‘overflow reservoir’ for fluid in cartilage. Previous studies indicate that chondrocyte metabolism is altered by fluid induced shear-stresses^30^ and that certain levels of fluid induced shear stress is healthy for chondrocytes^16^. Thus, fluid movement from cartilage to bone could also serve to mitigate possible cartilage tissue overload.

Fluid transfer between subchondral bone and cartilage could also participate in the cartilage degradation that follows joint injury. Recent studies demonstrate that the cartilage pericellular and extracellular matrices undergo significant alterations within three days after injury^7,9^. These changes may be caused, or exacerbated by, transport of cytokines and proteases from bone to cartilage. Changes to subchondral bone tissue after injury could affect fluid transport across the joint. Subchondral bone experiences a transient phase after joint injury of thinning and loss of bone mineral before becoming sclerotic^21,24^. These changes to bone geometry and density would be expected to impact how fluid is transported between the two tissues. Increased or decreased transport could each affect cartilage loading and nutrition. The impacts of fluid transport from physiological loading between cartilage and bone on the health of these tissues, and the role of altered fluid transport on deleterious cartilage changes following joint injury necessitates new directions of investigation.

Our model advances several aspects of the understanding of fluid transport between bone and cartilage. Using an ellipsoidal fiber distribution, we improved the modeling of collagen fibers within the cartilage tissue. This improvement is critical to osteochondral fluid transport models as it provides a more accurate structural representation of the mechanics in cartilage that are directly tied to this transport. Our model also improves the understanding of the fluid velocity vectors and fluid flux at the cartilage-bone interface during physiological loading by incorporating a biphasic viscoelastic ellipsoidal fiber distribution model for cartilage. This new approach improves cartilage mechanics modeling and thus fluid transport modeling for osteochondral studies. Lastly, validation of our model to established biphasic solutions improves our confidence about the material stress states underpinning the fluid transport between these tissues. The implications of fluid transport between cartilage and bone on the health of these tissues, as well as in degenerative joint diseases, are not yet understood. Several critical questions remain about how fluid transport between subchondral bone and cartilage affects chondrocyte health and viability, how transport between subchondral bone and cartilage following traumatic injury leads to increased inflammatory stimuli *(e.g.* TNFα) and how transport between subchondral bone and cartilage in healthy joints affect chondrocyte viability. These questions motivate new directions in investigating the role of fluid transport as a relevant factor in multi-tissue whole-joint health.

**Table 1.**

| Material | Elastic Modulus<br>E | Poisson's Ratio<br>$\nu$ | Porosity |
| --- | --- | --- | --- |
| Bone | 17 GPa | 0.290 | 5.5% |
| Superficial Zone Cartilage | $5.8539 - 5.7766Z + 4.0239Z^2 - 1.3551Z^3$ (MPa) | 0.499 | 73.0% |
| Middle Zone Cartilage | $5.8539 - 5.7766Z + 4.0239Z^2 - 1.3551Z^3$ (MPa) | 0.499 | 81.0% |
| Deep Zone Cartilage | $5.8539 - 5.7766Z + 4.0239Z^2 - 1.3551Z^3$ (MPa) | 0.499 | 86.0% |

## Supporting information

Supplemental Figure 1, Radial Mesh Convergence

Supplemental Figure 1, Axial Mesh Convergence

Supplemental Movie: model compression and stress relaxation

## Acknowledgements

We thank Dr. David Pierce for key insight and critical discussion in developing the model. Funding support provided by NSF (CMMI 1554708) and NIH (R01AR073964).

## Supplemental Figures

**Supplemental Figure 1:**
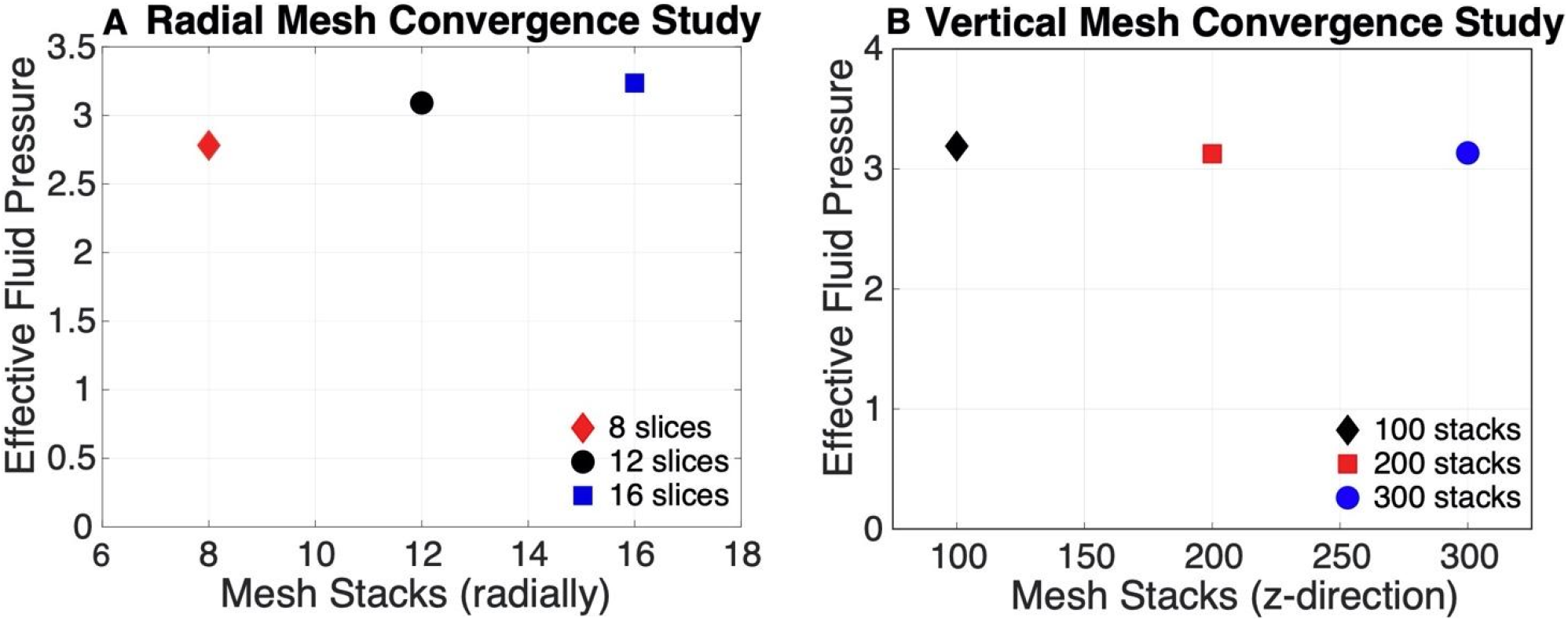
A: Radial mesh convergence study of the effective fluid pressure; B: Vertical mesh convergence study of the effective fluid pressure.

