## Supplementary figures and images for "Development and analytical validation of a finite element model of fluid transport through osteochondral tissue"

### Supplemental Figure 1, Axial Mesh Convergence

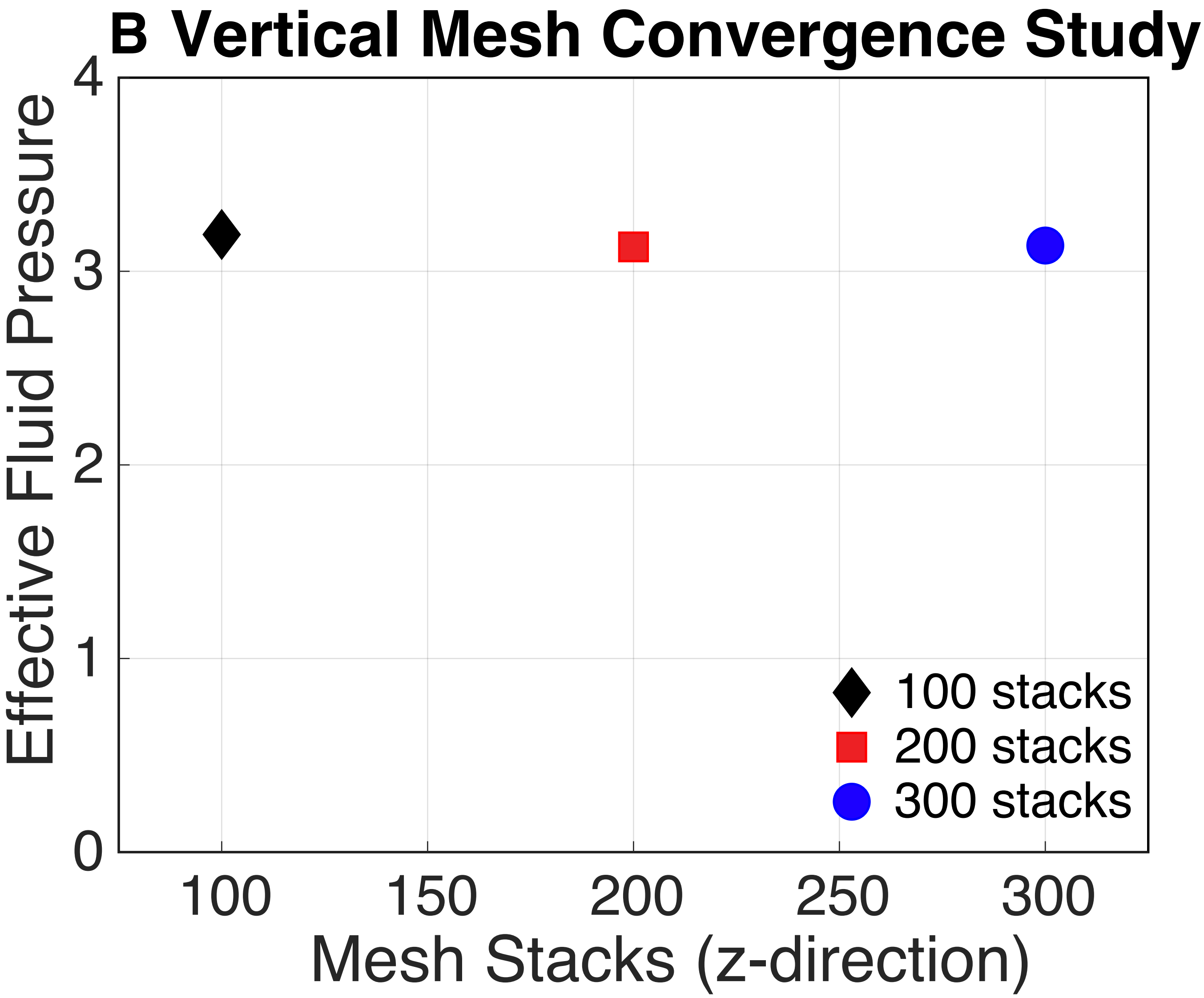

### Supplemental Figure 1, Radial Mesh Convergence

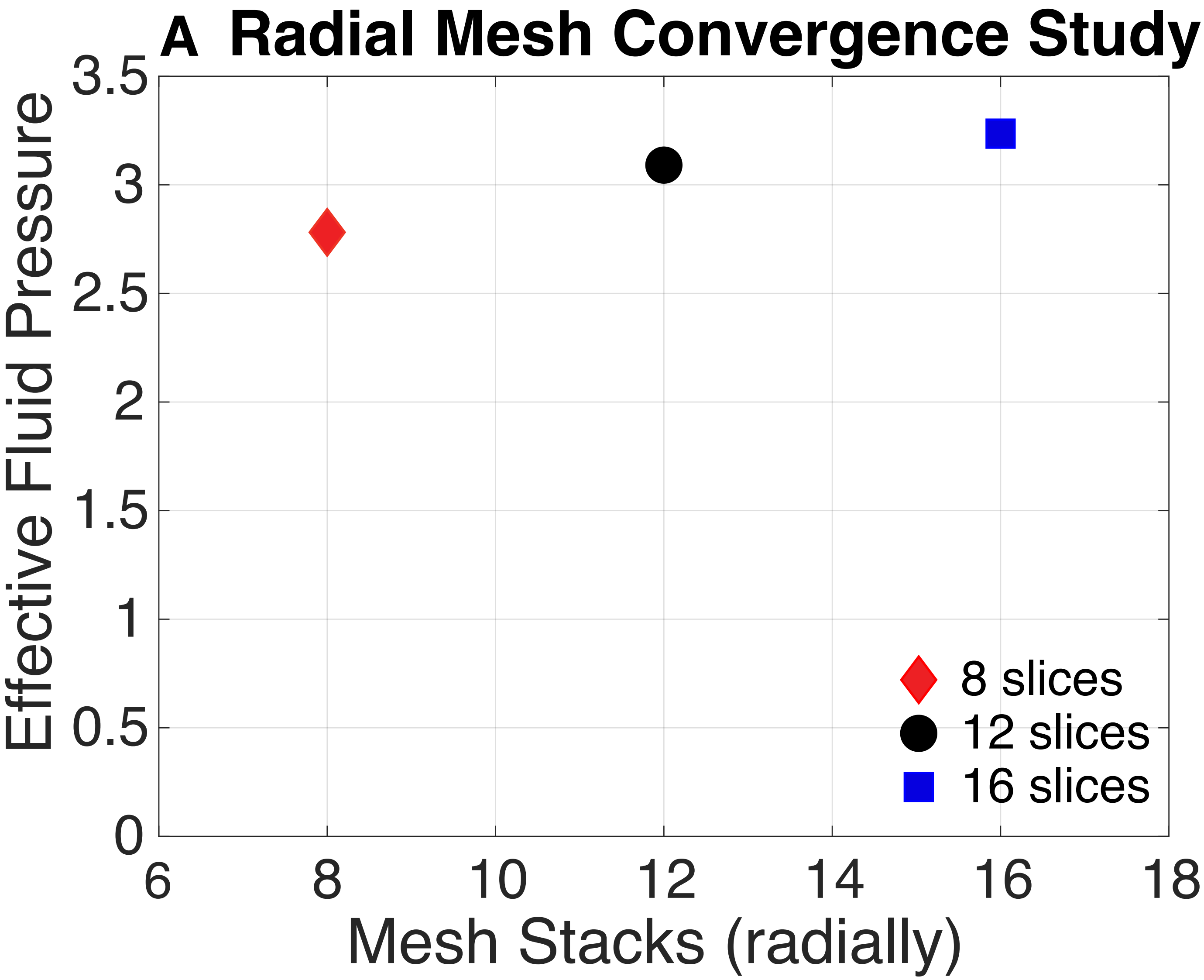
